# Free-flight kinematics of soldier flies during headwind gust perturbations

**DOI:** 10.64898/2026.03.31.715644

**Authors:** Dipendra Gupta, Sanjay P. Sane, Jaywant H. Arakeri

## Abstract

Large commercial and military aircraft can operate in a wide range of turbulent conditions, except during extreme weather events such as cyclones. Smaller man-made vehicles, such as micro aerial vehicles (MAVs) and nano aerial vehicles (NAVs), are significantly more sensitive to routine environmental wind fluctuations, making them difficult to control. In contrast, insects exhibit remarkable stability in naturally gusty conditions. Despite this, few studies have systematically investigated the impact of gusts and turbulence on insect flight performance. To address this gap and to gain fundamental insights into insect flight stability under gusty conditions, we examined the flight of freely flying black soldier flies subjected to a discrete head-on aerodynamic gust in a controlled laboratory environment. Flight motions were recorded using two high-speed cameras, and body and wing kinematics were analyzed across 14 distinct cases. In response to the gust, we observed consistent features across the cases: (1) asymmetry in wing stroke amplitude, (2) large changes in body roll angle—up to 160°—occurring over approximately two wing beats (∼20 ms) with recovery over ∼9 wing beats, (3) transient pitch-down attitude, and (4) deceleration in the flight direction. These rapid responses, combining passive and active control mechanisms, provide insight into the flight control strategies employed by insects. The findings offer valuable guidance for the design of MAVs and NAVs capable of robustly responding to gusts and unsteady airflow in natural environments.

## II. Introduction

The flying agility of insects had fascinated mankind from prehistoric times. Be it a windy or rainy day, they can forage and migrate successfully by maintaining stable flight. Atmospheric winds, however, offer a considerable challenge to these flying organisms to maintain the flight trajectory. The upper-speed limit of flight of a living or non-living flying organism is determined, in nature, by the combination of mean atmospheric wind speed and inherent turbulence of its environment. Larger commercial and military aircraft can fly in all except the most extreme weather conditions such as cyclones. In contrast, micro air vehicles (MAVs) and nano air vehicles (NAVs) are more vulnerable to environmental fluctuations than their bigger counterparts for both dynamics and environmental reasons. Dynamically, even a small change in wind speed and direction at a length scale comparable to MAVs can severely upset their flight trajectory and body orientation. Environmentally, MAVs are expected to operate at a lower altitude from the ground surface, full of turbulence and transient gusts. Maneuvering in a chaotic environment through the complex terrain of hills, canyons, trees, and street corners is, thus, a challenging task, but one that insects, birds, and bats routinely tackle. For the success of any operation, the autonomous controllability of MAVs using real-time onboard sensing of environmental gustiness is therefore crucial, raising the fundamental question of sensor and actuator placement on MAVs.

Despite turbulence and gustiness in the natural environment, the ability of small insects and birds to rapidly stabilize serve to inspire us to study their flight capabilities. Nonetheless, most studies focused on the aerodynamics of insects have been carried out on tethered, free-flying or dynamically scaled robotic model of insects [1, 2] and that too in still or laminar air in wind tunnels that are far from the real scenario insects diurnally experience. Very few studies have been carried out to understand the impact of gusts and turbulence on the flight performance of insects and birds [3]. These studies mostly demonstrate the effects of turbulence [4, 5] and a continuous trail of vortices [6-10] on the flight behaviour of insects with very few focused on isolated gusts [11]. Moreover, while turbulence and continuous vortex street are significant on a large scale, discrete gusts could be more relevant to smaller insects.

Prior studies have shown that severe wind conditions imposed on insects during flight can sharply upset their trajectory, flight speed, and body attitude. The insects manipulate, using passive, active, or a combination of both mechanisms, their body attitude to counteract the effect of these disturbances. One of the most intriguing features is that they respond to the perturbation on a very short time scale [12]. The characterization of the response as an active or passive mechanism is still not fully clear, possibly because of difficulty in the measurement of relevant kinematic or dynamic parameters. Selection of well-characterized gust that is relevant to insects, identification of the axis insects are highly susceptible to gust, and a clear distinction between active and passive mechanisms of stabilization can help us come up with a better and autonomous design for MAVs and NAVs.

In the present study, we explored the kinematic response of black soldier flies due to head-on gust. We perturbed insects with a controlled and well-characterized head-on vortex ring as a discrete gust [13, 14]. This method simulates the realistic environment that insects may face day-to-day during foraging and migration. Considering the head-on gust, we hypothesize that the forward speed of the flies will sharply decrease with the corresponding change in body pitch attitude. Further, given that insects are least stable along the roll axis, we conjecture that the response and recovery time along this axis will be shortest as well.

## III. Methods and materials

### A. Insect

We used soldier flies (*H. illucens*) maintained in a controlled laboratory environment. Their characteristic feature to fly towards light is one of the main reasons we chose them for study. This makes the control of their flight trajectory easier. For consistency across all the trial, we set certain criteria to select a fly before subjecting it to a gust-(a) No body parts (wings, halteres, etc.) were damaged, (b) Insect should be able to fly for at least 30 cm across the test section and (c) Its body length was minimum of 1cm and maximum of 1.5cm.

We released a group of 5-6 flies together into a test chamber using fly releasing duct (*FRD*) (to be discussed later in detail). Releasing in a group increases the probability of them being hit by the gust. After releasing, we would wait for the flies to fly themselves. We observed that all flies would, in some cases, fly at the same time just after release into the test chamber, while in other cases only one or none at all would fly into the test chamber. In latter cases, when an insect would remain stationary in *FRD* for more than 5 minutes, the lower internal wall of the duct was slightly hit with a plastic ruler to instigate its flight. However, while doing so, we ensured that flies weren’t hit directly, and no body parts were damaged. Some morphological and kinematic details of the flies are given in Table 1.

**Table 1.**
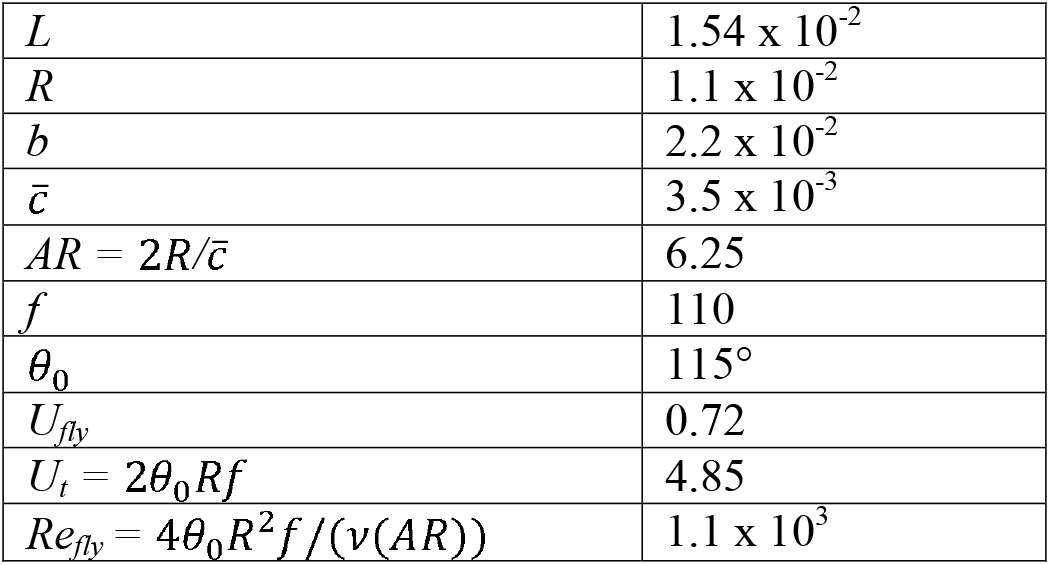
Morphological and kinematic details of soldier flies. All units are in *SI*.

### B. Gust perturbations

We used a loudspeaker to generate vortex ring as a gust perturbation (Fig.1). The speaker was enclosed in a wooden chamber (driving section) on the diaphragm side, to which a PVC nozzle of dimeter, *D*_0_ = 3.7cm was attached to facilitate the formation of the vortex ring. We generated a digital trapezoidal signal using NI-LabVIEW, converted it into analog form for physical output using NI–c9263, amplified it using an in-house DC power amplifier, and fed it to the speaker, resulting in the formation of a vortex ring at the exit of the nozzle.

We selected vortex ring velocity of 6.4 m/s to perturb the free-flying flies which is slightly higher than the wing-tip speed of the freely flying flies. We, however, also did some initial trials for gust velocity of 1-3 m/s but didn’t observe visual change in the body kinematics of the flies (not reported here). The vortex ring lasts at any location for a duration of approximately 1WB. For details of the gust generation and characterization, readers are requested to refer refs. [13, 14].

### C. Experimental setup and videography

We carried out the experiment in a closed test section made up of a Perspex chamber (Fig. 1). We used two synchronized high-speed cameras (Phantom VEO 640L and V611) to record the flight motion before, during, and after the gust. One camera captured the side view of the flight motion while the other an oblique top view. It is, however, noted the cameras weren’t placed orthogonal and that the non-orthogonality was taken care of by employing a rotation matrix that was determined using two orthogonal axes common to all the images. The cameras were operated at 4000 fps, 1280 x 800 resolution, and 70μs exposure time and were focused in the region of interest (*ROI*) where the speed of the gust becomes nearly invariant with space and time. We selected *ROI* (2*D*_*0*_ x 2*D*_*0*_ x 2*D*_*0*_) at a distance of 3*D*_*0*_ from the nozzle exit plane. Its volume was so chosen that the insect wouldn’t possibly go beyond the focal volume of the camera, even after its encounter with gust.

**Fig. 1.**
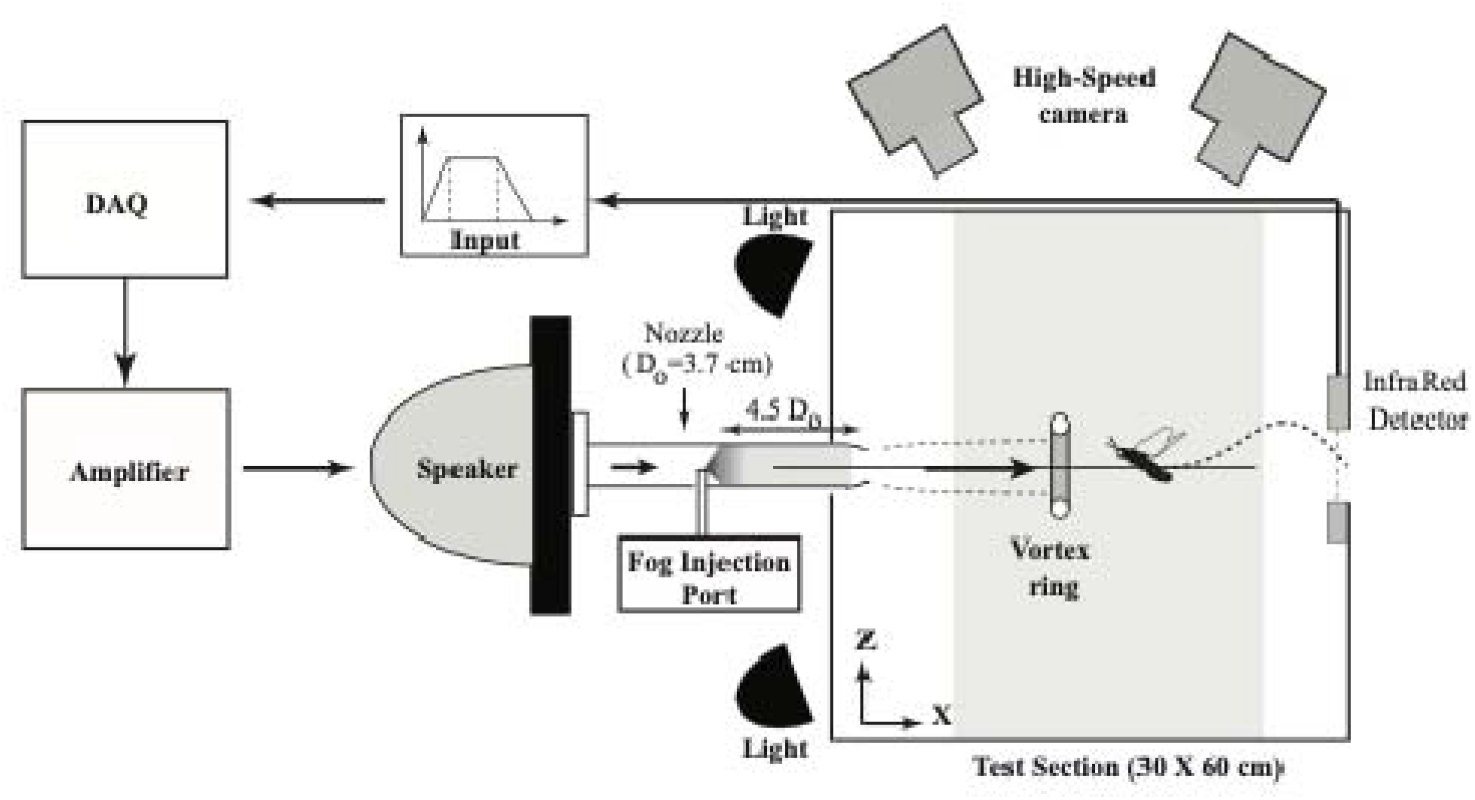
Gust generator system. Different components of the experimental setup and their arrangement. Test chamber, Speaker, nozzle, Infrared motion sensor, halogen lamp, two high-speed cameras, DAQ, and amplifier together constitute the gust generator system. Grey shaded region in test section is the region of interest where the encounter of insects with the gust is desired. Shown here is also the input signal which is converted into analog form by DAQ, amplified by an amplifier, and fed to the speaker for gust generation. All dimensions are in cm.

We placed an in-house infrared motion detector at the exit of *FRD* to initiate, based on a pre-specified time delay, the recording of the flight motion, and the generation of the gust. Detection of flight motion, automatic triggering of gust generator, and cameras were electronically synchronized.

### D. Kinematics reconstruction and analysis

We digitized six different body parts (head, abdomen, right and left-wing bases, and the corresponding wingtips) of the flies common to both the camera views using open-source MATLAB routine-DLTDV7 under manual supervision (Fig. 2) [15]. The DLT residual was kept less than 2 for each point and across videos. We used a glass capillary (75 mm long and 1mm diameter) as a calibration object (wand), one end of which we marked with black color for the distinction between its endpoints, and before any flight experiments, we moved it randomly throughout the calibration volume and recorded it. We then digitized the endpoints and used it as an aid to obtain DLT coefficient for image reconstruction using easywand5 [16].

**Fig. 2.**
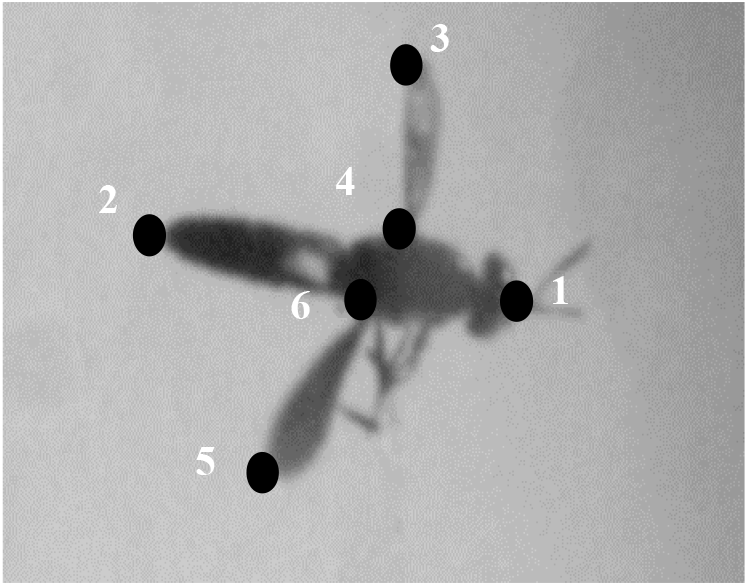
Kinematic extraction using recorded images. Oblique top view of fly showing digitization of different parts-head (1), abdomen (2), left wingtip (3), left wing base (4), right wingtip (5) and right wingbase (6).

We analyzed videos (700-1500 frames each) from 14 experiments using custom code written in MATLAB. We first down-sampled the image data to 1000 frames per second, and then, digitized each of the six different points on the body that was tracked to get their 3D position in the global reference frame. We assumed Centre of Mass (*CoM*) at one-third body length from the head towards the abdomen and used it to represent the flight trajectory. We defined X-axis as the direction along the nozzle centerline, and Y and Z lateral and vertical axes, respectively. To obtain flight velocity (*u,v*, and *w*) along X, Y, and Z directions, we first applied fourth-order low pass Butter-worth filter with cut-off frequency 200Hz to the corresponding *CoM* data and then calculate the velocity along each axis using second-order central difference scheme. We calculated total velocity (forward speed or velocity magnitude) as the resultant of three velocity components (i.e. 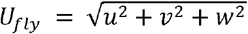, and non-dimensionalized each velocity component by the product of the body length of the flies (*L*) and their wingbeat frequency (*f*).

We calculated body pitch (*β*) as the elevation angle of a vector joining head and *CoM* of the body, relative to the horizontal plane. Similarly, the azimuthal angle of the same vector on the horizontal plane gives heading angle (*ψ*), and the elevation angle of the vector joining the wing bases gives body roll angle (*γ*) with respect to the horizontal plane. We subtracted the initial heading angle from the instantaneous heading angle to calculate the body yaw angle (*ψ*), since it would represent actual body yaw. Counter-clockwise rotation (i.e. rotation of insect from its right to left) in yaw and roll is considered positive, and pitch-up rotation is taken positive. We applied a fourth-order low pass Butter-worth filter with cut-off frequency 200Hz to the raw body angle data to smoothen out high-frequency noise that may have arisen due to digitization.

To find the stroke angle, we first converted the 3D position data of different digitized parts of the flies from a global coordinate system to a body reference system, beginning with a right-handed yaw rotation about the global vertical axis, followed by pitch and roll rotation about the global axis. The azimuthal angle of a vector joining wing tip and its base in stroke plane relative to the body reference system gives the measure of wing stroke angle (*θ*). We calculated the time period, and hence, the frequency of a wing beat from the videos by counting the number of frames in a wing beat. We calculated peak difference (*Δθ*_*0*_) as the absolute difference in stroke amplitude of right and left wings.

We chose to consider response and recovery time differently for each parameter-velocity, body angles, and wing stroke angle. For flight trajectory, velocity, and body angles, we defined response time as the time when the fly starts opposing the change induced by a gust on its flight motion, while recovery time as the time when the rate of the change becomes near-zero. Similarly, for wing stroke angle, the time when asymmetry in right- and left-wing stroke amplitude increases by more than ± 15° in response to the gust gives a measure of response time, while recovery time is defined as the time when asymmetry completely ceases or if persists is not more ± 15° for at least 3 successive wingbeats. We note that the definition of response and recovery time used here is somewhat different from that used in existing literature where changes in wing kinematics alone have been considered as the sole determinant of these time scales [6, 8, 11, 12, 17, 18]. One main advantage for defining response and recovery time for each kinematics differently is that it allows us to compare the response and recovery times for each parameter separately, and this can further help us discover if the fly responds to gust on a different time scale for each body and wing kinematic parameters and the role of wings (be it passive or active), if any, in stabilization definitively.

## IV. Results

### A. Trajectory and Flight speed

The flies were hit by the gust at an axial location of X_0_ = 4.16 ± 0.51D_0_ (mean ± SD) from the exit of the nozzle, and their lateral and vertical positions of *CoM* just before being hit by the gust were within the gust (Fig. 3A). In most cases, if the flies were to the right side of the gust, they continued to fly in the same direction even after being hit by the gust, except for cases 4, 14, and 10, where the opposite trend was observed. Similarly, they mostly flew downward after the encounter with gust in most of the cases except for 1,3,4,9 and 11 (Fig. 3B). The flies didn’t recover along Y and Z directions after gust in any of the cases.

**Fig. 3.**
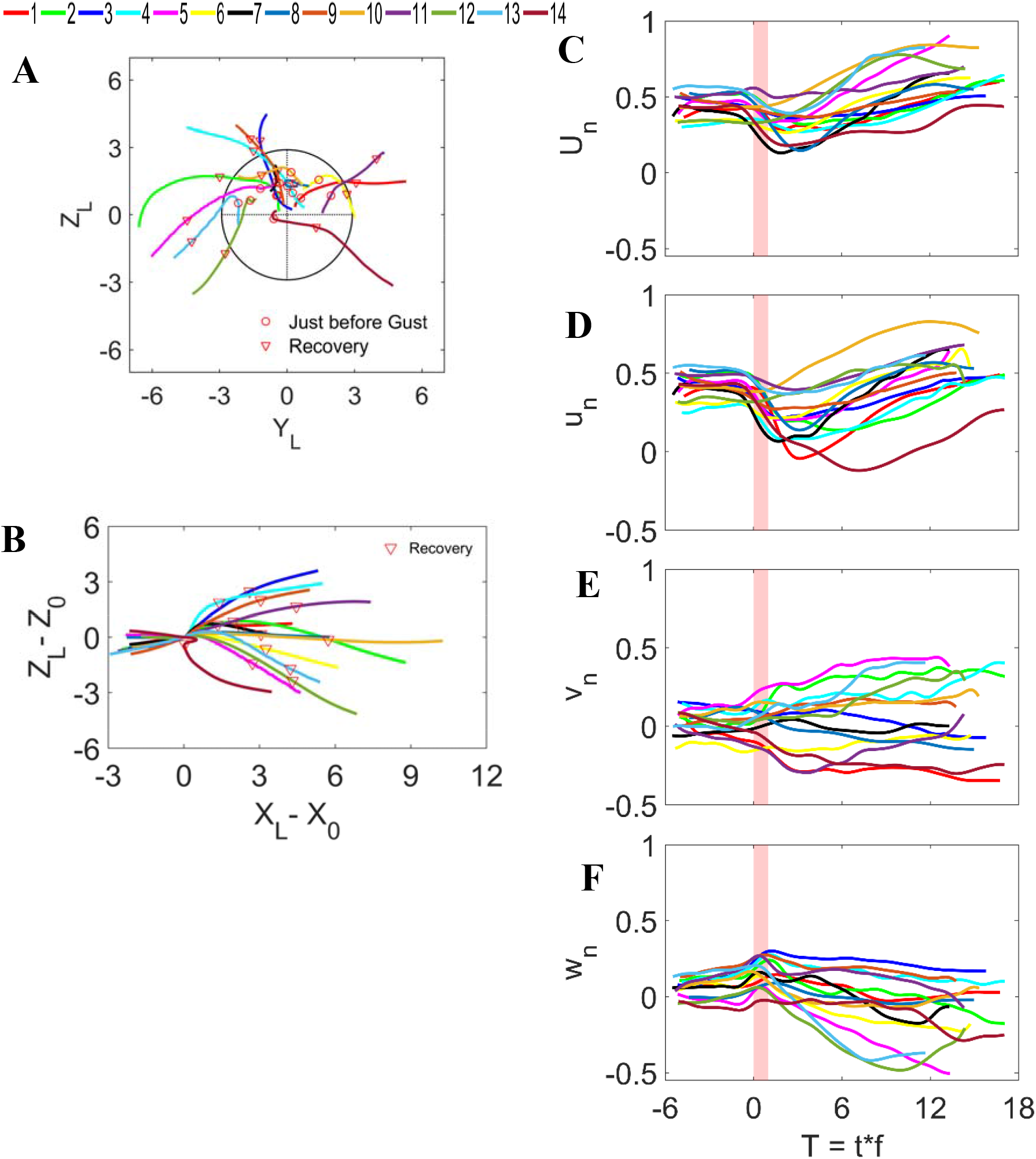
Flight path and velocities. (A) Trajectory in Y-Z plane normalized by the average body length (*L*) of flies. The mean axial distance where the gust hits the insects is *X*_*0*_ = 4.16 ± 0.51 *D*_*0*_. Black circle indicates the front view of the vortex ring and the intersection of vertical and horizontal dashed lines is the center of the vortex ring. Colored lines are the trajectories of flies for each trial. (B) Trajectory in X-Z plane, where (0, 0) indicates the position where the flies were hit by gust. (C-F) Non-dimensional total flight speed (*U*_*n*_), axial velocity (*u*_*n*_), lateral velocity (*v*_*n*_), and vertical velocity (*w*_*n*_) are plotted against the number of wingbeats (*T*). Velocities are non-dimensionalized by average wingbeat frequency and average body length, while the instantaneous time when multiplied by wingbeat frequency, gives the number of wingbeats. *T=0* indicates the time instance when the flies were just hit by the gust and the vertical pink shade the time during which flies were in contact with the gust.

Before being hit by the gust, the velocity of the flies was near-constant in each case and the dimensionless axial velocity of the fly was 0.36 ± 0.08 (mean ± SD), lateral velocity -0.03 ± 0.11 and vertical velocity 0.14 ± 0.09, indicating low velocity in transverse direction compared to the axial one (Fig. 3C-E). During the period of gust (∼ 1 wing beats), velocity along the axial direction decreased by 26%, while it increased by 53% and 4% on average along Y and Z directions. The overall velocity magnitude decreased by 15% on average during the gust.

During response time (∼ 3.5 wing beats after gust), axial velocity on average decreased by 36% while lateral velocity increased by 150% and vertical velocity decreased by 83%. The velocity magnitude during this period decreased on average by 30% in all cases, except for two cases 10 and 12 where the flight velocity increased by 90% and 135% respectively.

After recovery time (∼ 13 wing beats after gust), the axial, lateral, and velocity magnitude when compared to that before gust, on average, increased by 47%, 250%, and 55% respectively, and the vertical velocity decreased by 225%. The flies after recovery mostly maintained constant velocity except for cases 5 and 11 where they further accelerated.

Two main observations in the flight velocity are (a) sudden decrease in forward speed due to gust till response time and (b) velocity after recovery period higher than that before gust in all cases.

### B. Body and Flight angles

Before being hit by the gust, the initial orientation of the fly across the trials, along yaw, pitch and roll axes was 1 ± 37° (mean ± SD), 25 ± 13°, and 7 ± 10° respectively (Fig. 4A-C). Due to the gust, the absolute change (irrespective of the direction of rotation) in the roll was 83 ± 37°, with a maximum value of 161° counterclockwise (CCW). Similarly, body yaw changed by 29± 17°, with a maximum 69° CW, and pitch changed by 31 ± 14° with a maximum 61° pitched down.

**Fig. 4.**
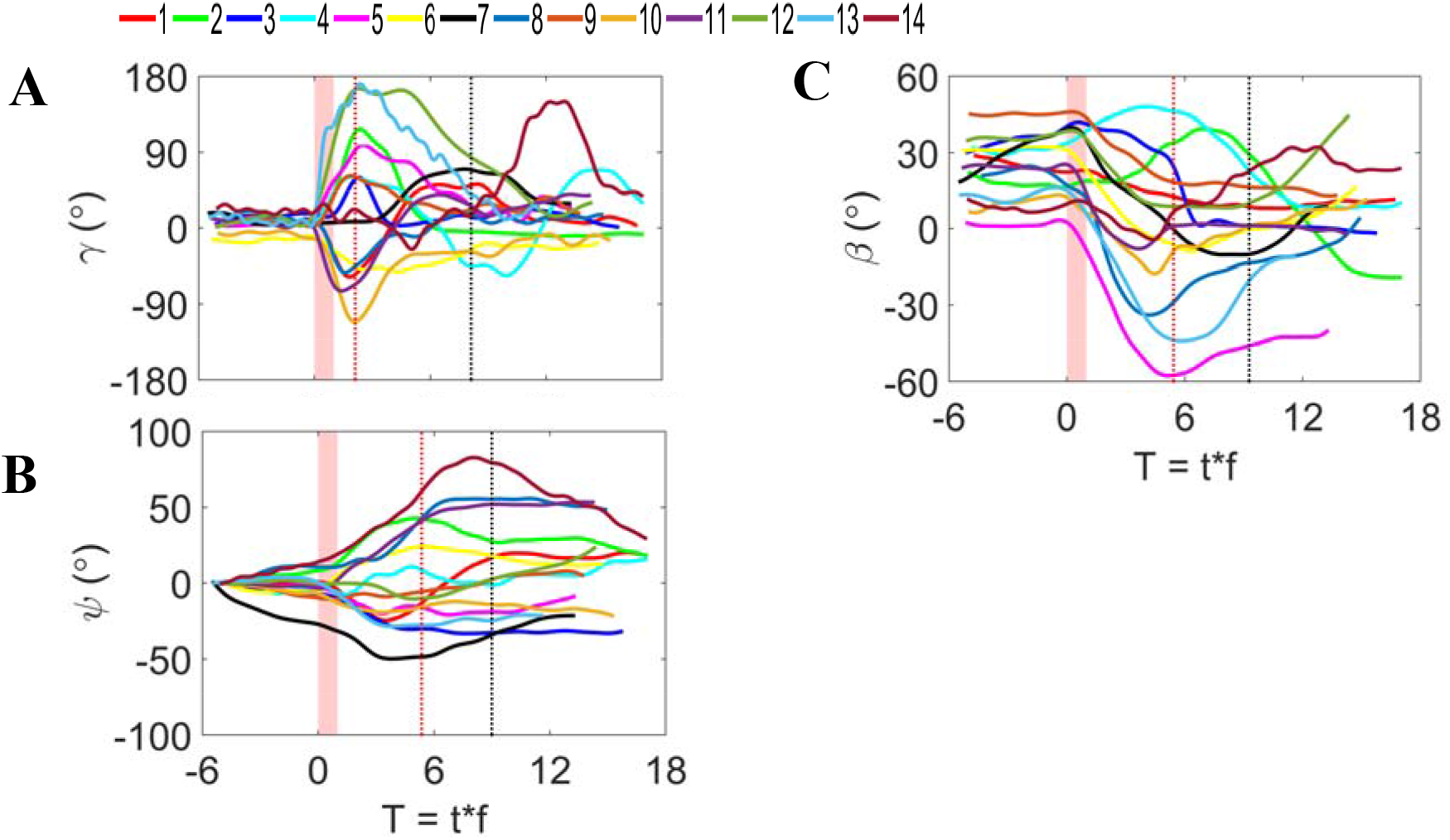
Body and flight angles and impact of gust on them. (A) Body roll (*γ*), (B) body yaw (*ψ*), and (C) body pitch (*β*), each plotted against the number of wingbeats. T=0, pink shade and color of each curve convey the same meaning as in Fig. 3. Red and black dashed lines indicate the mean response and recovery time respectively.

After recovery, the error calculated as the difference between the body orientation after recovery and that before gust was 11 ± 9° in roll with maximum 30° (CCW) (Fig. 4A-case 11), 23 ± 16° in yaw with maximum 56° (CCW) (Fig. 4B-case 14) and 25 ± 11° in pitch with maximum 47° pitched down (Fig. 4C-case 5).

### C. Wing Kinematics

Instantaneous wing stroke angle is shown for case 3 in Fig. 5A. Before being hit by the gust, the wing half stroke amplitude was observed to be 115 ± 17° (mean ± SD) and the peak difference between right and left wings was less than 15° for all cases (Fig. 5B, C). Results show that stroke asymmetry starts in 2.2 ± 1.5 wing beats (20 ±14 x 10^-3^ ms). The asymmetry continued for 8.9 ± 3 wing beats (81 ± 27 x 10^-3^ ms), after which either stroke became symmetric or asymmetricity if existed, was less than 15°.

**Fig. 5.**
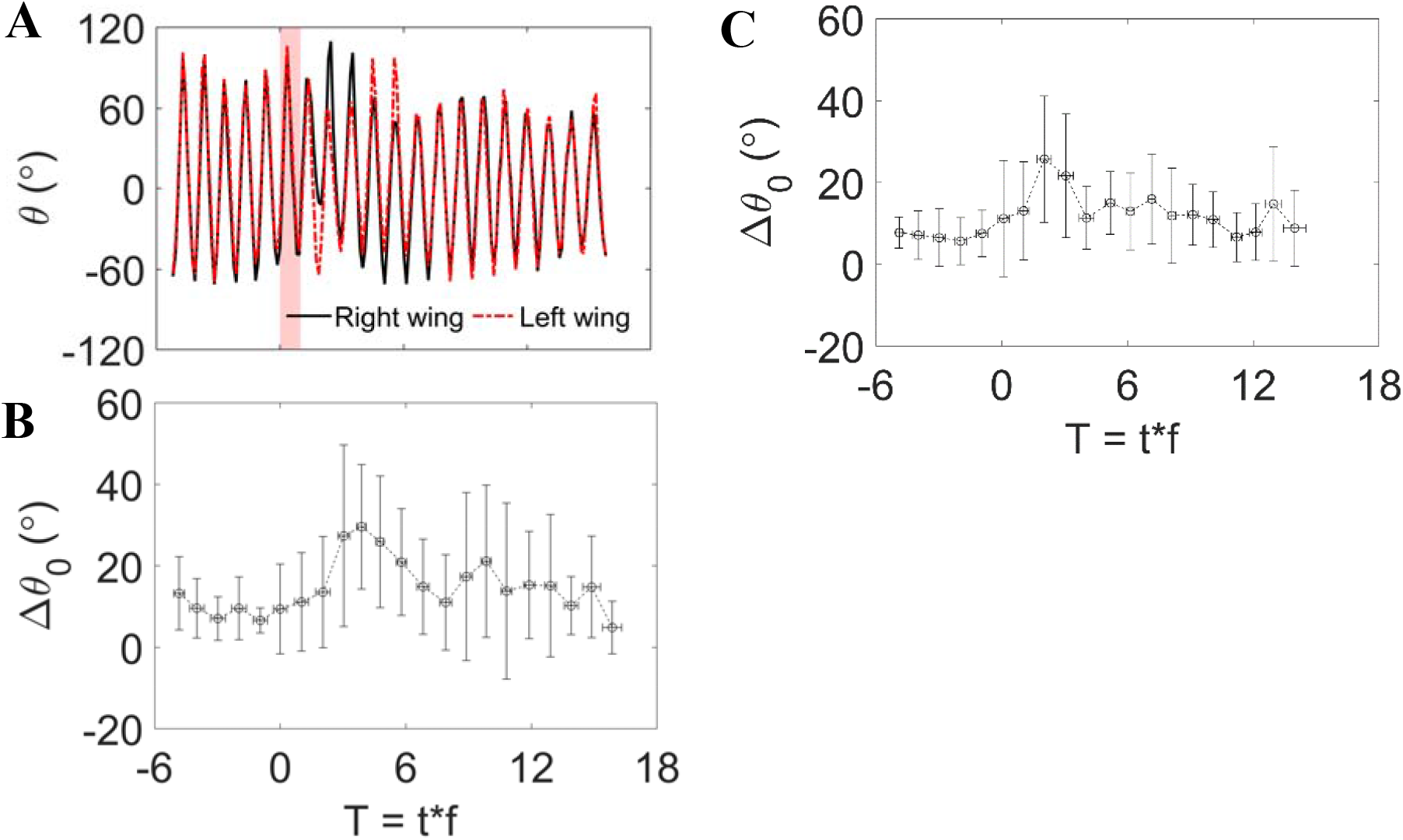
Wing stroke versus wingbeats. (A) shows a representative instantaneous wing stroke angle (θ) against wingbeats for case 3 (dark blue in Fig. 3 and Fig. 4). (B) and (C) show absolute values of difference in positive (dorsal) and negative (ventral) peak stroke amplitude (Δθ_0_) of right and left wings respectively. Values are absolute mean ± SD in both horizontal and vertical axes.

## V. Discussion

### A. Effect of gust on flight trajectory, speed and body angles

The diameter of the vortex ring was approximately 3.5 times the wingspan of the flies studied in this experiment. In all the cases, the initial position of the insect *CoM* was within the circumference of the vortex ring before being hit by gust. But, on the encounter with the gust, the flight trajectories altered in all three directions, and flies flew outside the circumference of the gust in both horizontal and vertical planes. Consequently, unlike bumblebees [10], none of the flies in our study recovered their lateral and vertical position after gust. Considering flight speed and the velocity of gust each of similar value in both the studies, the inability of soldier flies to recover their trajectory may be attributed to different perturbation methods (thin continuous air sheet in the latter case), body and wing morphology, flapping frequency and wing loading. One important observation is a continuous decrement in flight altitude in majority of the cases (Fig. 3B), which may be the result of decrement in vertical velocity by approximately 2 times after gust (Fig. 3F).

Similarly, like orchid bees, hawkmoths, and bumblebees, soldier flies also displayed lower forward speed immediately after gust [4, 6, 10, 19]. This implies possible increment in aerodynamic drag on the body due to gust and turbulence indicating that such severe environmental conditions possibly act as a limiting factor for flight performance [4]. However, we also observed that the constant forward speed of soldier flies after recovery, in all the cases, was higher than that during the normal flight before being hit by the gust. This suggests that, though the flies interact with gust for only 1WB, the effect of gust prevails longer. The higher speed after recovery may be attributed to the inertia associated with some active stabilizing mechanisms. It can also be a strategy to cope up with anticipated gusts.

Soldier flies, on average, displayed quite higher rotation along roll axis than that along pitch and yaw axes (Fig. 4), and response time was 2.1 ± 0.4 wing beats (WB), 5.4 ± 2.8 WB and 5.5 ± 1.3 WB along roll, yaw and pitch axes respectively (Fig. 6), indicating the least response time along roll axis, followed by yaw and pitch. Similarly, recovery time was noted 8.1 ± 2.6 WB, 9 ± 4.7 WB, and 9.3 ± 3.4 WB along roll, yaw and pitch axes respectively (Fig. 6), indicating the flies take approximately same time (∼ 9 WB) to correct the body orientation along the three axes. Response time along roll axis is on average 2.5 times faster than that along pitch and yaw axes and agrees well with the findings of ref. [12]. Beatus et al. (2015) also compared the roll response time in fruit fly with existing literature on yaw [17] and pitch [18] correction in gust and found that it was 3.5 times faster than yaw response and 2.5 times than pitch response. In contrast, soldier flies responded to gusts along both yaw and pitch axes on the approximately same time scale. Also, we note that the flies took a longer time (13.13 ± 4.27 WB) to recover the flight speed compared to body and wing angles. This aberration may be due to the inertia of motion even after stabilization in body angles.

**Fig. 6.**
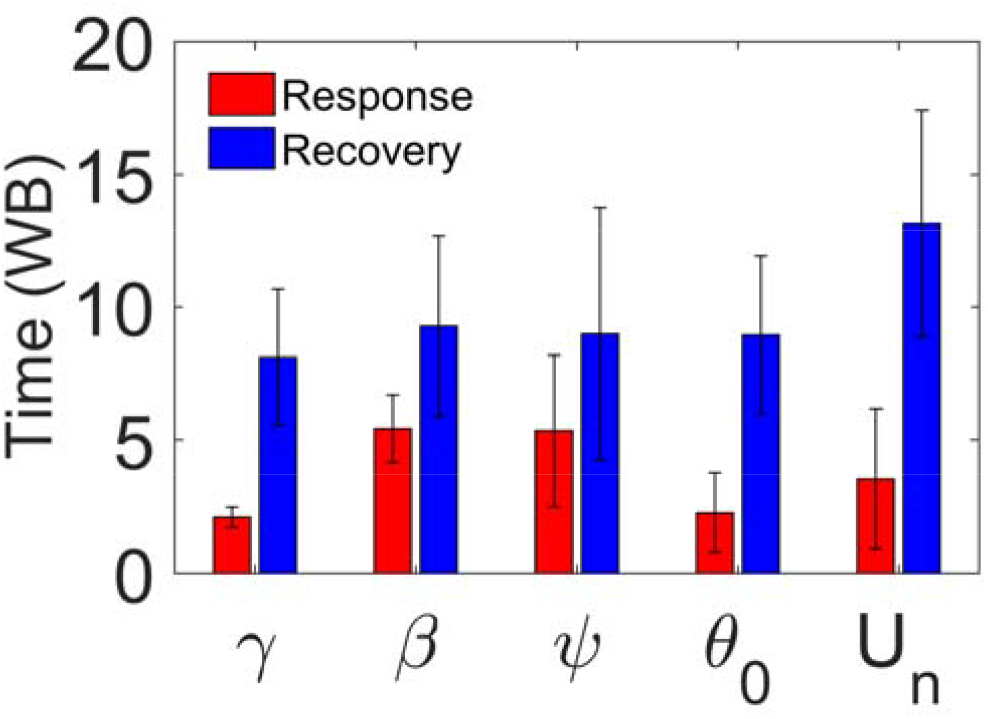
Response and recovery time for body and wing kinematics. *γ, β, ψ, θ*_*0*_ and *U*_*n*_ denote body roll, pitch, yaw, wing-stroke amplitude and non-dimensional flight speed. Response time of roll and wing stroke amplitude, and yaw and pitch are approximately same, and the case is same for recovery time of body angles and stroke amplitude. Recovery in flight speed takes longer than the body angles and stroke amplitude. Values are mean ± SD.

Response to gust along roll axis on such a short time scale across all trials when compared to that along pitch and yaw axes shows that the flies are highly susceptible to gusts along the roll axis and more immune to disturbance along the other two axes. This may be attributed to the lowest moment of inertia along roll axes which, for the same amount of torque, induces a large change along the roll axis than that along other axes [4, 12, 20]. Further, the flies recovered to near-zero roll orientation after recovery time in all the cases. However, unlike in roll, they never attain the initial pitch and yaw orientations, but mostly stabilize to new body orientations exhibiting neutral stability along these axes, suggesting that zero roll is the most preferred orientation.

Similarly, the highest angular velocity was found along the roll axis, followed by that along the pitch and yaw axes. This observation, at least along roll axes, is consistent with the findings of refs. [11, 12], despite the employability of different methods of perturbation. In particular, soldier fly in case 13 interestingly rolled by more than 160° in less than 3 wingbeats. What is even more intriguing is that despite a quite higher angular rate (∼28000°/s) for this case, the fly stabilizes in less than 9WB. The observations regarding pitch and yaw rates are, however, in contrast to that in stalk-eyed flies studied by ref. [11], where the authors reported yaw angular rate higher than pitch angular rates by ∼3 times, while the present study shows that yaw and pitch average angular rates are approximately same.

Such a short response time, high angular rates, and short recovery time along each axis suggest that passive mechanism alone cannot be accounted for these observations. The flies must also be employing some active mechanism to recover from perturbation caused by the gust [10, 11, 17, 18, 21].

### B. Possible active mechanisms for stabilization

Apart from employing passive mechanisms like large body inertia for gust mitigation, insects have been observed to actively respond to perturbations by altering their wing and body kinematics. Some insects extend their hind or forelimbs on the encounter with gust to bring stability along the roll axis [4, 7, 10, 12, 22]. Similar observation is also noted in present study in soldier flies (Fig. 7). In most cases, before gust, all the limbs were tucked against the body or against each other. The limbs, in the latter case, appeared as if hanging vertically from the body. After gust, not only hind limbs or forelimbs but all of these limbs are pulled upward and stretched laterally outward with respect to the body in all the cases. Spreading of limbs is associated with an increase in rotational moment of inertia aiding to attenuate the effect of gust on body roll [4]. Even after stability, the limbs remained spread-out, though lesser than that during recovery.

**Fig. 7.**
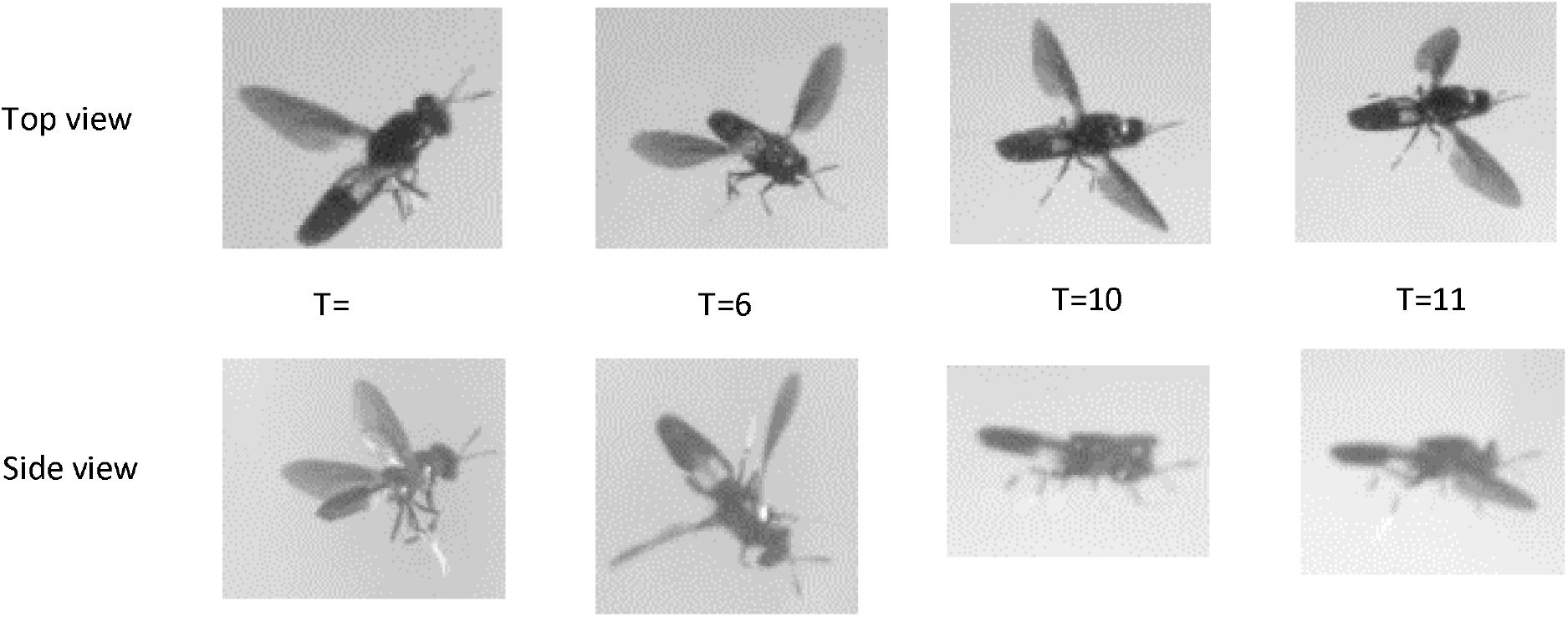
Limb deflection in soldier fly due to gust. Limbs are pulled upward and spread laterally outward upon hit by gust. T indicates time in wing beats.

Further, asymmetry in wing stroke amplitude starts on a time-scale similar to the response time along roll axis (Fig. 6), and it is interesting to note that the flies recovered the body attitude (roll, yaw and pitch angles) and wing stroke symmetricity, on average, after the same time period (∼ 9WB) (Fig. 5B, C and Fig. 6). Moreover, we did not observe any change in the wingbeat frequency of soldier flies due to gust. So, considering the time latency for any passive response in a fly (> 5–6 WB), beginning of stroke asymmetry on such a short timescale (∼2 wing beats) is believed to be an active mechanism employed by insects (flies and bees) to correct for perturbations [4, 11, 12, 17-19, 23]. Existing literature also reports that insects employ differential angle of attack to correct for yaw in response to gusts [11, 17, 19] and also employ differential wing excursion forward or backward on each wing to correct for pitch [19, 23]. The present study, however, due to lack of this information, does not permit us to comment on the role of angle of attack and wing excursion on stability. We note that abdominal deflection, as in some insects, for pitch stability was not observed in the present study [6, 22, 24] (see Fig. 7).

## VI. Conclusion

We investigated how black soldier flies respond to a discrete head-on aerodynamic gust using a vortex ring in a confined chamber. Flies pitch down and decelerate upon gust encounter and show extreme sensitivity along the roll axis, with maximum excursions of 160° at angular rates of ∼28,000°/s. Remarkably, they recover near-zero roll within ∼9 wing beats, while pitch and yaw adopt new sustained orientations, indicating neutral stability in these axes. Wing kinematics reveal that stroke asymmetry initiates within ∼2 wing beats and returns to symmetry after ∼9 wing beats, closely matching the recovery of body orientation. Roll corrections coincide with stroke asymmetry onset, whereas pitch and yaw corrections are ∼2.5 times slower, demonstrating that asymmetric wing motion is a primary mechanism for rapid gust stabilization. These fast responses suggest a combination of passive and active control strategies. While this study focused on kinematics, the aerodynamic forces underlying stabilization remain unquantified. Future work could employ volumetric PIV with freely flying insects or use robotic insect models with force measurements and planar PIV to directly link wing aerodynamics with gust response.

## I. Nomenclature

*AR*: wing aspect ratio
*b*: wingspan
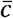: wing mean chord length
*CoM*: center of mass
*D*_0_: nozzle exit diameter
*f*: wing beat frequency
*FRD*: fly releasing duct
*fps*: frame per second
*L*: average body length
*R*: wing radius
*Re*_*fly*_: flight Reynolds number
*T*: Number of wingbeats
*U*_*fly*_: flight average velocity
*U*_*t*_: wingtip velocity
*u,v,w*: X, Y, and Z components of flight speed
*U*_*n*_: non-dimensional flight velocities
*WB*: wingbeat
*θ*_0_: peak-to-peak stroke amplitude
*ν*: kinematic viscosity

## Author Contributions

D.G., S.P.S. and J.H.A. designed the project. D.G designed and constructed the experimental setup, carried out the experiments, processed and analyzed the data, and wrote the manuscript. J.H.A. and S.P.S. supervised the project.

## Acknowledgments

Funding for this study was provided by grants from the Air Force Office of Scientific Research (AFOSR) # FA2386-11-1-4057 and # FA9550-16-1-0155, and National Centre for Biological Sciences (Tata Institute of Fundamental Research) to SPS. We also acknowledge the support of the Ministry of Earth Sciences, Government of India, under project no. MESO-0034 and the Department of Atomic Energy, Government of India, under project no. 12-R&D-TFR-5.04-0800.

## Notes

### Competing Interest Statement

The authors have declared no competing interest.

